# What’s under the Christmas tree? Soil acidification alters fir tree rhizosphere bacterial and eukaryotic communities, their interactions, and functional traits

**DOI:** 10.1101/2021.03.16.435746

**Authors:** Blaire Steven, Jacquelyn C. La Reau, Stephen J. Taerum, Nubia Zuverza-Mena, Richard S. Cowles

## Abstract

pH has been identified as a master regulator of the soil environment, controlling the solubility and availability of nutrients. As such, soil pH exerts a strong influence on indigenous microbial communities. In this study we describe a soil acidification experiment and the resulting effects on the rhizosphere communities of fir trees on a Christmas tree plantation. The acidification treatment reduced the pH of bulk soil by ∼1.4 pH units and was associated with reduced Ca, Mg, and organic matter content. Similarly, root chemistry differed due to soil acidification with roots in acidified soils showing significantly higher Al, Mn, and Zn content and reduced levels of B and Ca. 16S rRNA and 18S rRNA gene sequencing was pursued to characterize the bacterial/archaeal and eukaryotic communities in the rhizosphere soils. The acidification treatment induced dramatic and significant changes in the microbial populations, with thousands of 16S RNA gene sequence variants and hundreds of 18S rRNA gene variants being significantly different in relative abundance between the treatments. Additionally, co-occurrence networks showed that bacterial and eukaryotic interactions, network topology, and hub taxa were significantly different when constructed from the control and acidified soil rRNA gene amplicon libraries. Finally, metagenome sequencing showed that the taxonomic shifts in the community resulted in alterations to the functional traits of the dominant community members. Several biochemical pathways related to sulfur and nitrogen cycling distinguished the metagenomes generated from the control and acidified soils, demonstrating the myriad of effects soils acidification induces to rhizosphere microbes.

**IMPORTANCE:** Soil pH has been identified as the property that exerts the largest influence on soil microbial populations. We employed a soil acidification experiment to investigate the effect of lowering soil pH on the bacterial and eukaryotic populations in the rhizosphere of Christmas trees. Acidification of the soils drove alterations of fir tree root chemistry and large shifts in the taxonomic and functional composition of the communities, involving pathways in sulfur and nitrogen cycling. These data demonstrate that soil pH influences are manifest across all organisms inhabiting the soil, from the host plant to the microorganisms inhabiting the rhizosphere soils. Thus, pH is an important factor that needs to be considered when investigating soil and plant health, the status of the soil microbiome, and terrestrial nutrient cycling.

## INTRODUCTION

Current estimates suggest that ∼40% of the world’s arable soils are acidic (pH 6.0 or below (1)) and their extent is expanding (2). Broadly reported consequences of soil acidification are: decreases in plant species richness, lower plant productivity, loss of soil organic carbon stocks, leaching of nutrients from the soil, and increased N_2_O flux (2–5). In agricultural soils, soil acidification is generally driven by the application of ammonia, urea, and elemental sulfur fertilizers, or growth of certain crops such as legumes which can acidify soils (6–8). Soil pH is a driver of multiple soil characteristics. For instance, soil acidification is linked to lower concentrations of organic matter, changes in the quality of soil organic matter, and the solubility and concentration of ions contained in soils (9–11). As such, soil pH is an essential metric in determining the health and functionality of soils.

Soil pH has been identified as one of the primary soil characteristics that influence the diversity and composition of the indigenous soil microbial communities (12–15). Yet, many of these studies have been performed at the continental scale, or at sites with a pH gradient, and thus pH is only one of multiple edaphic factors that differ among sampled locations (9, 15). In general, studies that investigate the interaction between soil pH and soil microbiology have focused on bulk soil samples and not the root associated soils that constitute the rhizosphere. As plant roots absorb and exchange ions with the soil, the pH at the root surface can often be 1-2 pH units different than surrounding soils (16, 17). For example, when plants absorb NO_3_^-^ they raise the pH whereas utilizing NH_4_^+^ lowers pH (16). Thus, plants influence the local pH of the rhizosphere and may magnify or dampen bulk soil pH changes for plant associated microorganisms (18).

In this study we describe the results of a soil manipulation study in a Christmas tree farm planted with Canaan fir (*Abies balsamea* (19)). Rhizosphere samples were collected approximately six years after the initial acidification treatment. We endeavored to test if the acidification effect was still present, the influence of the acidification on the nutrient status of the roots, and the composition of the rhizosphere archaeal, bacterial, and eukaryotic communities through sequencing of 16S rRNA and 18S rRNA genes. We additionally investigated the interactions of the rhizosphere communities through co-occurrence networks and the functional potential of the communities through metagenomic sequencing. Through these efforts we show that a history of soil acidification induces significant changes in the fir tree root tissue chemistry as well as the composition and functional potential of the rhizosphere microbial populations.

## MATERIALS AND METHODS

### Field site description

Christmas trees of the species *Abies balsamea* (L.) Mill. var. *phanerolepis* Fernald (Canaan fir) were located in Allen Hill Farm in Brooklyn, Connecticut (41.7696, −71.9183). Plots were 3.4 by 11.7 m planted with 14 trees at a spacing of 1.7 m between trees with a similar spacing of 1.7 m between rows. On June 18, 2014 pelletized sulfur (containing 90% sulfur) was applied to the plots at a rate of 3,370 kg per hectare. The sulfur was incorporated into the soil manually with a rototiller to a depth of 15 cm. In the soil, elemental sulfur is oxidized to sulfuric acid, reducing the pH (8). Three-year old 30 cm tall root transplants were planted in the field on April 13 and 14^th^, 2015. Soil pH was measured in August of 2015 and was 4.1 and 5.9 for the acidified and control soils, respectively. Samples for this study were collected on June 2^nd^, 2020, almost six years after the soil acidification was initially performed.

### Soil chemistry

Soil cores were collected to a depth of 10 cm from interspaces of trees in the same row. Three cores were collected per row and were composited into a single sample. Three replicate rows were sampled for both the control and acidified soils, resulting in three independent replicates for soil chemical analysis. Soil chemistry was performed with the Ag Soil test at Spectrum Analytic (https://www.spectrumanalytic.com/) using standard methods.

### Root collection

Fine roots were uncovered with an ethanol-sterilized trowel in the vicinity of an individual tree. Rhizosphere samples were processed in a manner similar to that described by McPherson *et al*. (20). Briefly, six *∼*10 cm sections of fir tree fine roots were collected from each tree and shaken vigorously to remove loosely adhering soil. The root sample was then transferred to a 50 ml plastic centrifuge tube containing 25 ml of sterile phosphate buffered saline (PBS) (Ingredients of PBS by weight in 1 liter of deionized water were: 8 g of NaCl, 0.2 g of KCl, 1.44 g of Na_2_HPO_4_, and 0.24 g of KH_2_PO_4_). The samples were then stored on ice until further processing in the field. Roots were collected from four individual trees per row, with three rows sampled per soil treatment, resulting in 12 rhizosphere/root samples per soil treatment. The rhizosphere soil was removed from the roots in the field by vortexing the root samples for 2 minutes at full speed. The roots were then removed with ethanol-sterilized forceps and transferred to a sterile plastic sample bag. The soil remaining in the centrifuge tube after vortexing was considered the rhizosphere sample and was immediately stored on dry ice. Root and rhizosphere samples were transported to the lab in New Haven, Connecticut. The rhizosphere samples were stored at −80 °C, while the root samples were stored at −20 °C until further processing.

### Root nutrient analysis

To analyze elemental content in the roots, tissues were oven-dried at 65 °C for 48 h. About 0.2 g per dried sample was digested with 4 ml of concentrated nitric acid (∼ 70%) for 45 min at room temperature, and for an additional 45 min at 115 °C in a hot block (DigiPREP System; SCP Science, Champlain, NY). After cooling, the final volume was fixed with deionized water. The elemental profile of the digests was determined by inductively coupled plasma optical emission spectroscopy (ICP-OES; iCAP 6500, Thermo Fisher Scientific, Waltham, MA). Yttrium was employed as an internal standard and a multi-element sample of known concentrations within the run for quality control purposes.

### Rhizosphere DNA extraction

The frozen rhizosphere samples were removed from the −80 °C freezer and rapidly thawed in a 55°C water bath. Soil particles and cells were collected by centrifugation at 14,000 rpm for 10 min at 4°C. DNA was extracted from the 0.25 g of soil from the resulting pellet using the DNeasy PowerSoil Kit (Qiagen). DNA extractions were verified by gel electrophoresis in a 1% agar gel.

### Amplification of 16S rRNA and 18S rRNA genes

Bacteria and archaea 16S rRNA genes were amplified with the 515F (GTGYCAGCMGCCGCGGTAA) and 806R (GGACTACNVGGGTWTCTAAT) primer pair (21). Extracts were each amplified with 10 µL Platinum SuperFi II DNA Polymerase (Invitrogen), which also included 7.5 µM of both the mPNA and pPNA peptide nucleic acid (PNA) clamps (mPNA: GGCAAGTGTTCTTCGGA and pPNA: GGCTCAACCCTGGACAG) to block amplification of host plant mitochondria and plastid rRNA genes, respectively (22). PCR conditions consisted of 94 °C for 2 min followed by 30 cycles of 94 °C for 15 s, 60°C for 15 s, 68 °C for 15 sec, and 4 °C for infinite hold. The resulting amplification products were verified by gel electrophoresis and cleaning and normalization of individual PCR products was performed with SequalPrep™ Normalization Plate (96) Kit (Invitrogen). The normalized PCR amplicons were mixed, and the quantity and quality of the DNA pool was verified using an Agilent TapeStation. The resulting 16S rRNA gene amplicons were submitted to the Yale Center for Genome Analysis for sequencing on the Illumina MiSeq platform using 2×250 bp chemistry.

For eukaryotic 18S rRNA gene amplification we first undertook to design a method to block the amplification of 18S rRNA genes from the host tree. A new peptide nucleic acid (PNA) clamp was designed based on the pipeline described in Taerum *et al.* (23). To obtain a reference 18S rRNA sequence for PNA clamp design, DNA was extracted from *A. balsamea* needles using a GeneJET Plant Genomic DNA purification Mini Kit (Thermo Scientific), following the manufacturer’s directions for DNA purification from lignified polyphenol-rich plant tissues. A 170 bp fragment of the 18S rRNA gene was amplified with the primers Euk1391F (5’-GTACA CACCGCCCGTC-3’) and EukBr (5’-TGATCCTTCTG CAGGTTCACCTAC-3’) (24). This fragment includes the V9 hypervariable region, which is one of the most frequently targeted regions for high throughput sequencing of eukaryotes. PCR was performed as described in Taerum *et al*., 2020 (23). The fragment was sequenced at The Keck DNA Sequencing Facility at Yale on an Applied Biosystems 3730xL DNA Analyzer.

PNA clamp design consisted of *in silico* fragmentation of the *A. balsamea* V9 sequence into 15-17 bp k-mers, which were then mapped to the SILVA database containing animal, plant and protist sequences (25). K-mers that did not match any animal, plant or protists sequences were then screened using PNA TOOL (https://www.pnabio.com/support/PNA_Tool.htm) to ensure the clamp had a melting temperature between 76 and 82°C and consisted of fewer than 35% guanines and 50% purines. The selected clamp (AbiesV9_01, with the sequence GTTCGCCGTCTTCGACG) was synthesized by PNA Bio, Inc. (Newbury Park, CA, U.S.A.).

Quantitative PCR was used to test the effectiveness of the clamp at different concentrations. Reactions consisted of 1 x SsoAdvanced Universal SYBR Green Supermix (Bio-Rad, Hercules, CA, U.S.A.), 0.5 µM of each primer, and 2 ng template for a total reaction volume of 10 mL. AbiesV9_01 was added to the reactions at a range of concentrations (0, 0.75, 1.5, 3.75, and 7.5 µM), with each reaction concentration being conducted in triplicate. Reactions were conducted on a CFX96 Touch Real-Time PCR machine (Bio-Rad) Thermal cycler and consisted of an initial denaturing step of 95°C for 2 minutes, followed by 40 cycles of 95°C for 10 s and 60°C for 15 s. A concentration of 7.5 µM was selected as it suppressed amplification of host 18S rRNA genes and matched the concentration used for 16S rRNA gene amplification.

Rhizosphere 18S rRNA genes were amplified with the primer pair 1391F and EukBr with Platinum SuperFi II DNA Polymerase (Invitrogen) along with the 7.5 µM of the AbiesV9_01 PNA clamp. The reaction conditions were 95 °C for 2 min, followed by 30 cycles of 95 °C for 15 s, 78 °C for 10 s, 60 °C for 30 s, and 72 °C for 30 s, and 4 °C for an infinite hold. The 18S rRNA amplicons were cleaned and normalized identically as for the 16S rRNA gene amplicons and submitted to the Yale Center for Genome Analysis for sequencing on the Illumnina MiSeq platform using 2×250 bp chemistry.

### Amplicon sequence analysis

Both 16S rRNA and 18S rRNA gene sequences were initially processed using the mothur software package (v. 1.44.2(26)). Quality filtering consisted of generating contigs and selecting for sequences of at least 253 b.p. in length for 16S rRNA genes and 80 b.p. in length for the 18S rRNA gene datasets. Chimeric sequences were identified with the VSEARCH algorithm (27) as implemented in mothur, using the most abundant sequences as a reference for chimera detection. All putative chimeric sequences were removed from the datasets. The 16S rRNA were classified against the SILVA v132 reference database using the RDP naïve Bayesian classifier (29) as implemented in mothur, and sequences identified as belonging to chloroplasts were removed (25). 18S rRNA gene sequences were classified with the PR^2^ database, also using the RDP naïve Bayesian classifier, and sequences identified as unclassified Eukaryotes were removed (28). The resulting set of sequences in both datasets were assigned to Amplicon Sequence Variants (ASVs) employing a 100% sequence similarity threshold.

The mothur output files were imported into the phyloseq R package for descriptive and statistical analyses (30). Prior to alpha-diversity calculations and NMDS ordinations the sequence datasets were subsampled (random without replacement) to the size of the smallest dataset to maintain equal sampling between datasets. To identify statistically significant differences in phylum taxonomic bins and ASV relative abundance unnormalized ASV count data was employed. Rare ASVs, consisting of 5 or less sequences, present in less than 20% of the samples were removed. Data were normalized using centered log-ratio transformations and statistically significant differences were identified with the ALDEX2 package (31).

### Network analysis

Co-occurrence networks were analyzed on ASVs consisting of at least 50 sequences for both the 16S rRNA and 18S rRNA gene sequence datasets. This resulted in a combined dataset of 396 ASVs (317 16S rRNA and 79 18S rRNA). Networks were analyzed with the NetCoMi package in the R software suite (32) employing the SPIEC-EASI metric for network construction (33). Associations were estimated with the SPRING approach (34) with the default normalization and zero handling settings. The nlambda and replication numbers were set to 100 and 20, respectively.

### Shotgun metagenomic sequencing

DNA samples from individual trees in the same row (i.e. 4 trees) were composited at equal molar ratio to produce templates for metagenome sequencing. In this manner, three replicate samples were sequenced for each soil treatment. Sequencing libraries were prepared using the Ligation Sequencing Kit (SQK-LSK109; Oxford Nanopore) and individually barcoded with the Native barcoding Expansion (EXP-NBD104; Oxford Nanopore). Libraries were sequenced for 72 hours on the Oxford Nanopore MinIon with the MinION Flow Cell (R9.4.1). Basecalling was performed with the Guppy software (4.2.2) using the accurate basecalling model. The resulting Fastq files were assembled with the Flye assembly software (2.8-b1674), using the metagenome settings (35, 36). A round of assembly polishing was performed with Racon v. 1.4.19 (37) followed by a second round of polishing with Medaka (v. 1.2.3 https://nanoporetech.github.io/medaka/) using default parameters. The resulting contigs were binned with metaBAT2 (38) and the resulting assembly bins were assessed with CheckM for completeness and contamination (39). Genes were identified and translated with the CheckM implementation of Prodigal (40) and the resulting amino acid sequences were assigned to KEGG pathways with the GHOSTX webserver (41–43)

## RESULTS

### Soil Chemistry

In 2020, six years after the initial sulfur treatment the pH of the acidified soils remained approximately 1.4 pH units lower than the control soils (Table 1). Soil acidification was also associated with significantly lower calcium, magnesium, and organic matter content than the control soils (Table 1). Thus, the effects of the pH treatment were still apparent during the current study and influenced several other soil parameters.

**Table 1.** Bulk soil chemistry.

| Plot | pH | Ca (ppm) | Mg (ppm) | P (ppm) | K (ppm) | OM (%) | CEC (mEq/100 g) |
| --- | --- | --- | --- | --- | --- | --- | --- |
| Acid 1 | 5.1 | 641 | 54 | 560 | 98 | 3.6 | 11.4 |
| Acid 2 | 4.9 | 401 | 40 | 470 | 78 | 3.8 | 12.8 |
| Acid 3 | 5.2 | 816 | 54 | 567 | 88 | 3.9 | 12.0 |
| Mean | 5.1 | 619 | 49 | 532 | 88 | 3.8 | 12.1 |
| Control 1 | 6.4 | 1,725 | 87 | 444 | 103 | 4.7 | 9.7 |
| Control 2 | 6.5 | 2,039 | 85 | 517 | 76 | 4.1 | 10.8 |
| Control 3 | 6.6 | 2,228 | 91 | 575 | 103 | 4.6 | 11.6 |
| Mean | 6.5 | 1,997 | 88 | 512 | 94 | 4.5 | 10.7 |
| P-value | 0.001 | 0.01 | 0.01 | ns | ns | 0.05 | ns |

pH, mineral availability (determined by Mehlich-3 extraction), organic matter and cation exchange capacity of soil from the root zone of Canaan fir trees. Each value represents soil combined from four trees per plot. Statistical significances between acidified and control soil determined by non-paired t-test on non-transformed values.

### Root tissue analysis

The mineral nutrition status of the root tissue was analyzed by ICP-OES. Three elements, B, Ca, and Na were present at significantly lower concentrations in roots from the acidified soils (36%, 47%, and 31% lower, respectively) whereas Al, and Zn were significantly more abundant in the roots from the acidified soils (56% and 47% higher; Figure 1). Note that prior to the ICP analysis the roots were rinsed with PBS to remove rhizosphere soil. This procedure may have influenced the measured values for Na, P, and K, and so the values we report for these elements should be considered relative rather than absolute values in the root tissue for comparisons between the roots in control and acidified soils. Yet, these data clearly demonstrate that the acidification treatment of the soils translated into altered chemistry of the fir tree root tissue.

**Figure 1.**
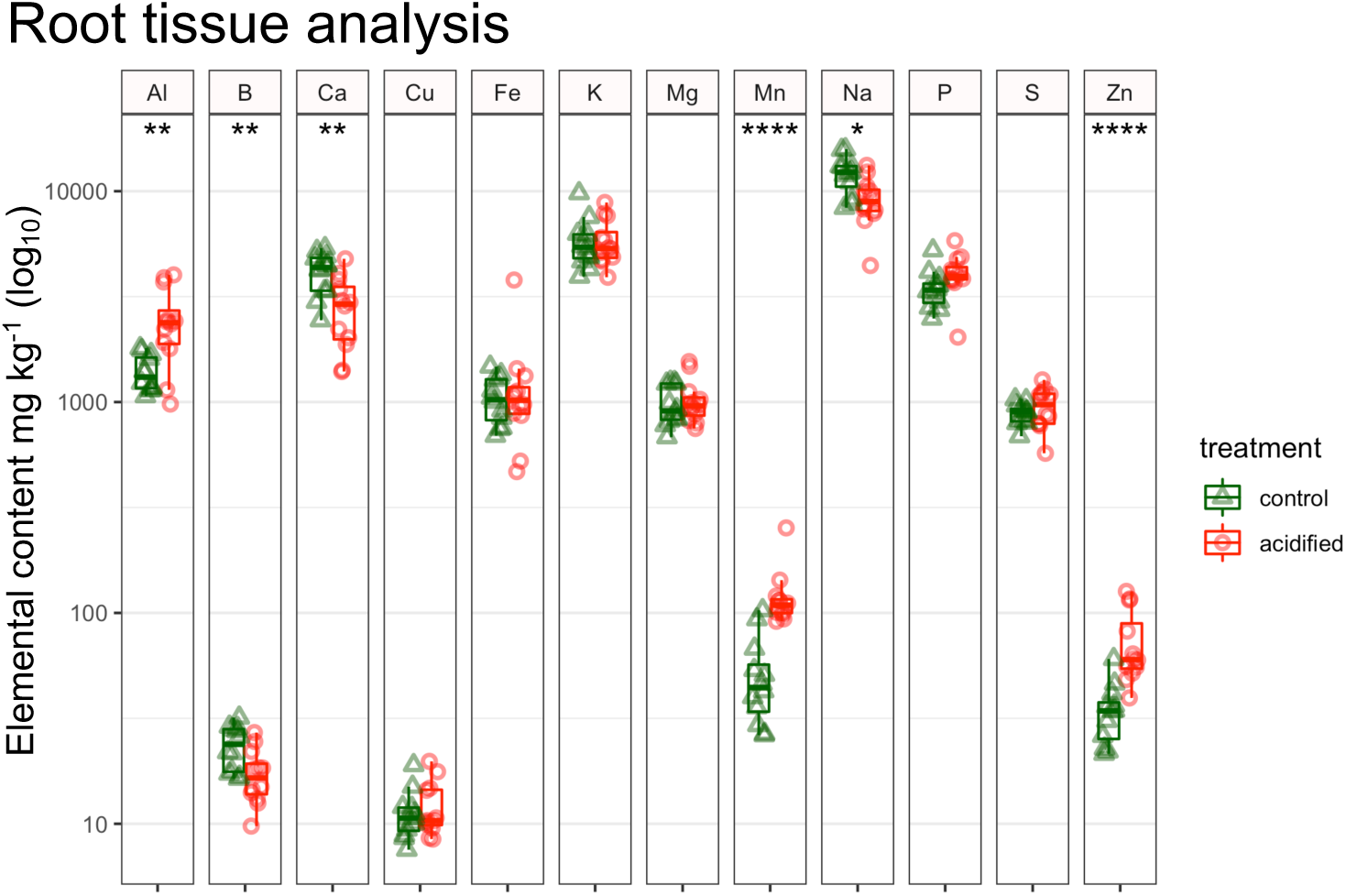
ICP-OES analysis of root tissue. Note that the roots were washed in a solution of phosphate buffered saline, which explains the high Na, P, and K values (see methods for formula). Thus, these values should only be considered relative to the acidification treatment. For both panels the significance of a t-test is denoted by the asterisks, which represent * P <= 0.05, ** P<=0.01, *** P<=0.001, **** P<=0.0001.

### Diversity of 16S rRNA and 18S rRNA gene libraries

Sequencing of the 16S rRNA gene was pursued to investigate the Bacterial and Archaeal populations in the fir tree rhizosphere whereas 18S rRNA gene sequencing was used to characterize the Eukaryotic populations. The alpha diversity of the 16S and 18S rRNA gene libraries was assessed to determine if the acidification treatment influenced community diversity of the fir tree rhizosphere. A statistically higher number of 16S rRNA gene amplicon sequence variants (ASVs) were recovered from roots in the control soils (mean 44,830 ASVs) compared to the acidified soils (39,868 ASVs), an 11% decrease in the number of recovered ASVs (Figure 2A). Similarly, the Shannon’s Diversity Index of the 16S rRNA gene libraries was significantly higher in the control soils (Figure 2B). For the 18S rRNA gene libraries, a mean of 4,266 ASVs were recovered from the control rhizospheres versus a mean of 3,734 ASVs from roots in the acidified soils (Figure 2A), a 12% decrease in association with the acidification treatment. Yet, for the 18S rRNA gene libraries the Shannon’s Diversity Index was similar between the control and acidified samples (Figure 2B). Taken together, these data suggest that the 16S rRNA gene datasets were more diverse than the 18S rRNA gene datasets, pointing to a more diverse bacterial/archaeal community in comparison to their eukaryotic counterparts. Additionally, the acidification treatment was associated with a significant decrease in the number of recovered ASVs for both the 16S and 18S rRNA gene libraries, suggesting that soil acidification resulted in a trend towards decreased diversity of both the 16S rRNA and 18S rRNA gene amplicon datasets.

**Figure 2.**
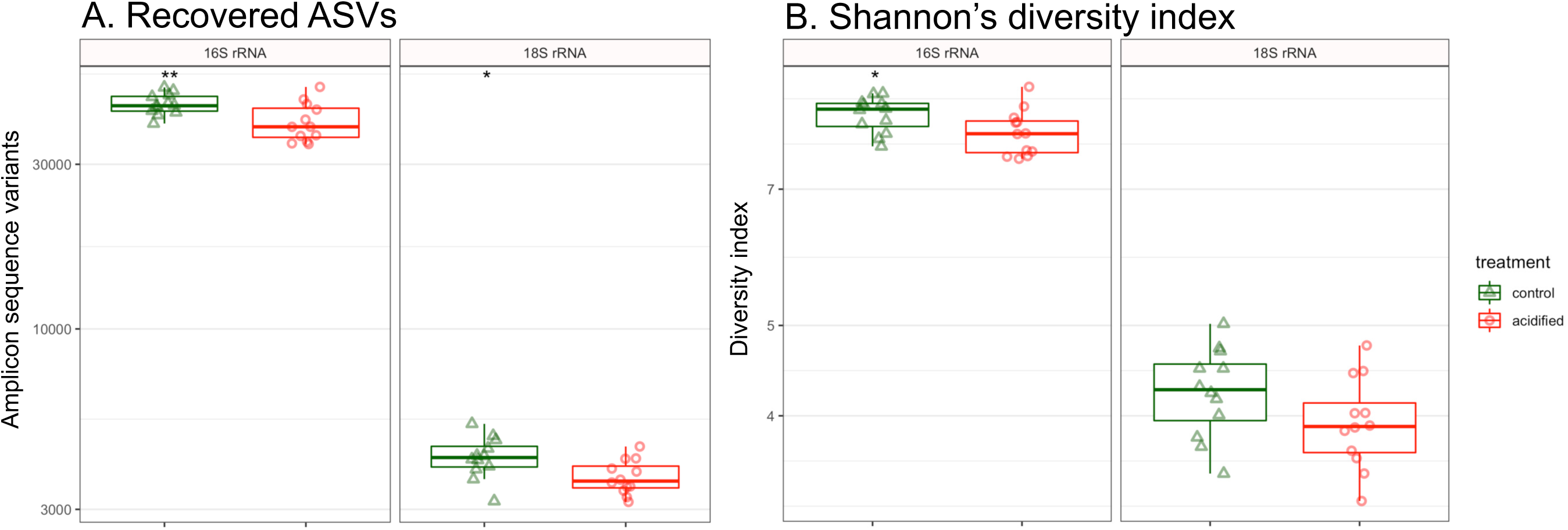
Alpha diversity of 16S rRNA and 18S rRNA gene datasets. **A. N**umber of recovered ASVs. **B.** Shannon’s diversity index based on ASV relative abundance. For both panels the significance of a t-test is denoted by the asterisks, which represent * P <= 0.05, ** P<=0.01.

### Alterations in 16S rRNA libraries due to soil acidification

The relationship between sequence datasets was visualized with non-metric multidimensional scaling (NMDS) and showed that the control datasets clearly clustered distinctly from the acidified samples, with a P-value ≤ 0.001 (Figure 3A). Communities for each tree with acidified soil had greater inter-sample distances in the NMDS plot, suggesting that soil acidification inflated community heterogeneity amongst individual rhizosphere samples. This was confirmed by comparing the inter-sample dissimilarity between samples, which was larger for the datasets from acidified soil and was highly significant (P ≤ 0.0001; Figure S1).

**Figure 3.**
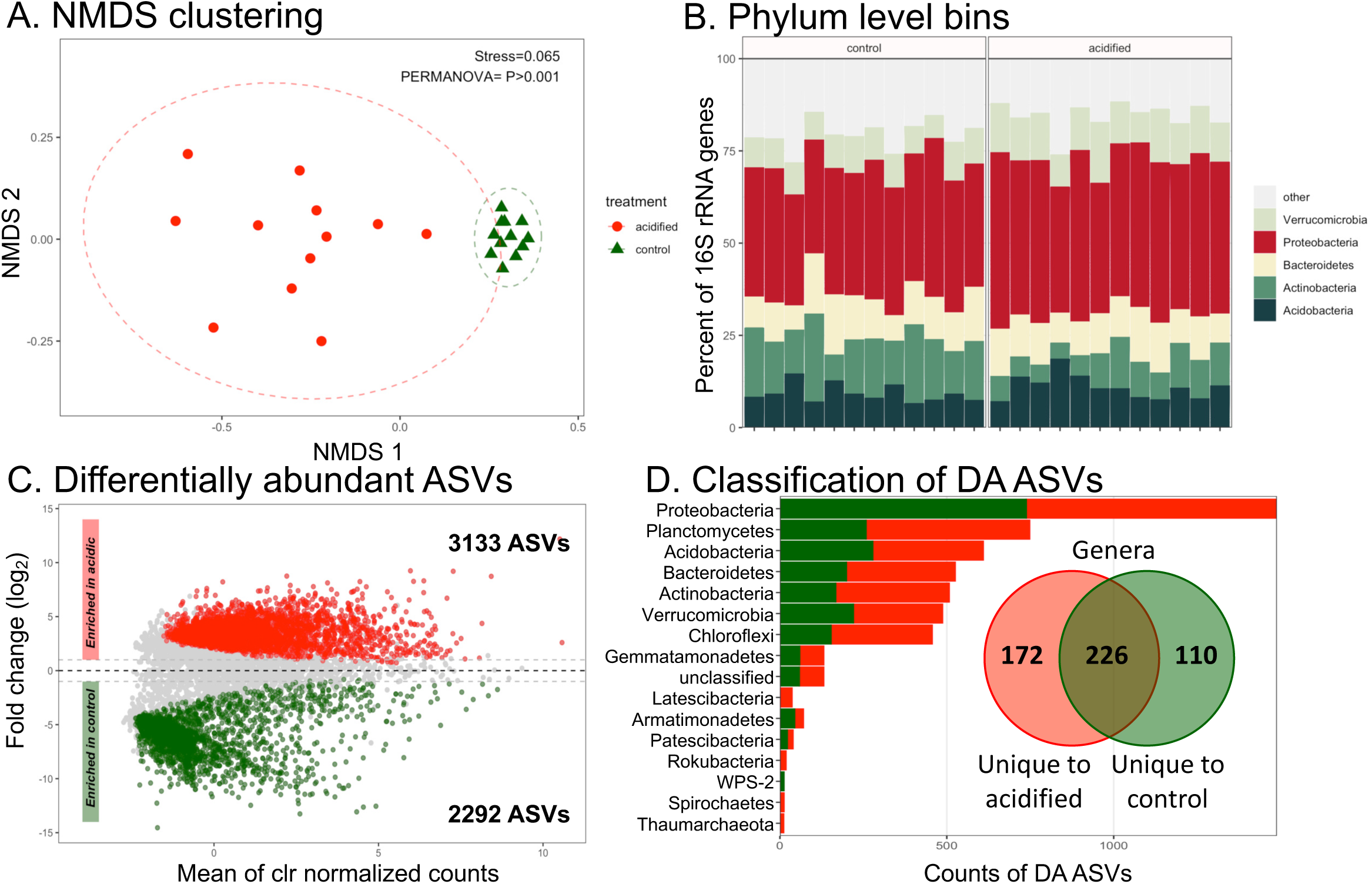
Composition of 16S rRNA gene datasets. **A.** Nonmetric multidimensional scaling ordination on ASV level data. The stress value of the ordination and P-value of a PERMANOVA statistical test are indicated. Ellipses denote 95% confidence intervals fitted onto the ordination. **B.** Phylum-level taxonomic bins in the datasets. Phyla consistently accounting for greater than 1% of sequence reads are displayed with the remainder assigned to the category “other”. A table of phyla showing statistically significant differences in relative abundance are shown in Table S1. **C.** MA plot displaying differentially abundant (DA) ASVs. The number of DA ASVs enriched in each condition are indicated in the inset text. **D.** Taxonomic classification of DA ASVs. Each bar represents the sum of DA ASVs classified to each Phylum. The inset Venn diagram shows ASVs classified to the genus level. The diagram shows the sum of genus level bins uniquely enriched in the control or acidified soils, with the overlap indicating genera with members enriched under both conditions.

The 16S rRNA gene sequences were classified to the phylum level. A total of 37 phyla were identified in the dataset, with the five most abundant phyla generally making up >75% of sequence reads (Figure 3B). Overall, the relative abundance of phyla was similar between the control and acidified rhizospheres. Yet, 20 phylum level bins were identified as significantly different in relative abundance due to the pH treatment (Table S1). For instance, an increase of Protobacteria was associated with acidification, rising from 35% of the control sequence libraries to 42% in the pH treatments. Taken together these data show that the pH treatment was associated with shifts in the taxonomic composition of the rhizosphere communities at broad taxonomic levels.

We additionally tested for significant differences in ASV relative abundance due to soil acidification. Thousands of ASVs were found to be significantly different in response to the pH treatment (Figure 3C). The differentially abundant (DA) ASVs belonged to 15 different phyla (Figure 3C). Most of the DA ASVs belonged to the phylum Proteobacteria matching their dominance in the datasets (Figure 3B). However, the second largest class of DA ASVs were the Planctomycetes, which were among the rare “other” phyla in the rhizosphere (Figure 3B, 3D). Yet, Proteobacteria and Planctomycete-related ASVs were identified as being enriched in both the control and acidified soils, suggesting that there was not a consistent response across the groups. There were some DA ASVs that belonged to phyla specifically enriched in the acidified soils, namely the Latescibacteria, Rokubacteria, Spirochetes, and the archaeal phylum Thaumarchaeota. Similarly, WPS-2 related DA ASVs were unique to the control soils (Figure 4D). These observations are likely influenced by the relative rareness of these taxa in the datasets (Figure 3B) and thus may not be a true reflection that these taxa are particularly sensitive to the acidification treatment. The DA ASVs could be further classified to 508 different genus level bins (Figure 4C, inset Venn diagram). An interesting observation was that a large proportion of the differentially abundant genus-level bins (226 or 44%) contained genera with ASVs that were identified as being more abundant in both the control and acidified soils. This suggests that much of the response to acidification was occurring at a sub-genus level, *i.e.,* between species of the same genus, or even specific ASVs. Taken together, these data show that a diverse set of taxa were identified as responding to the soil acidification treatment and few taxa showed a particular sensitivity to soil acidification in one direction or the other.

**Figure 4.**
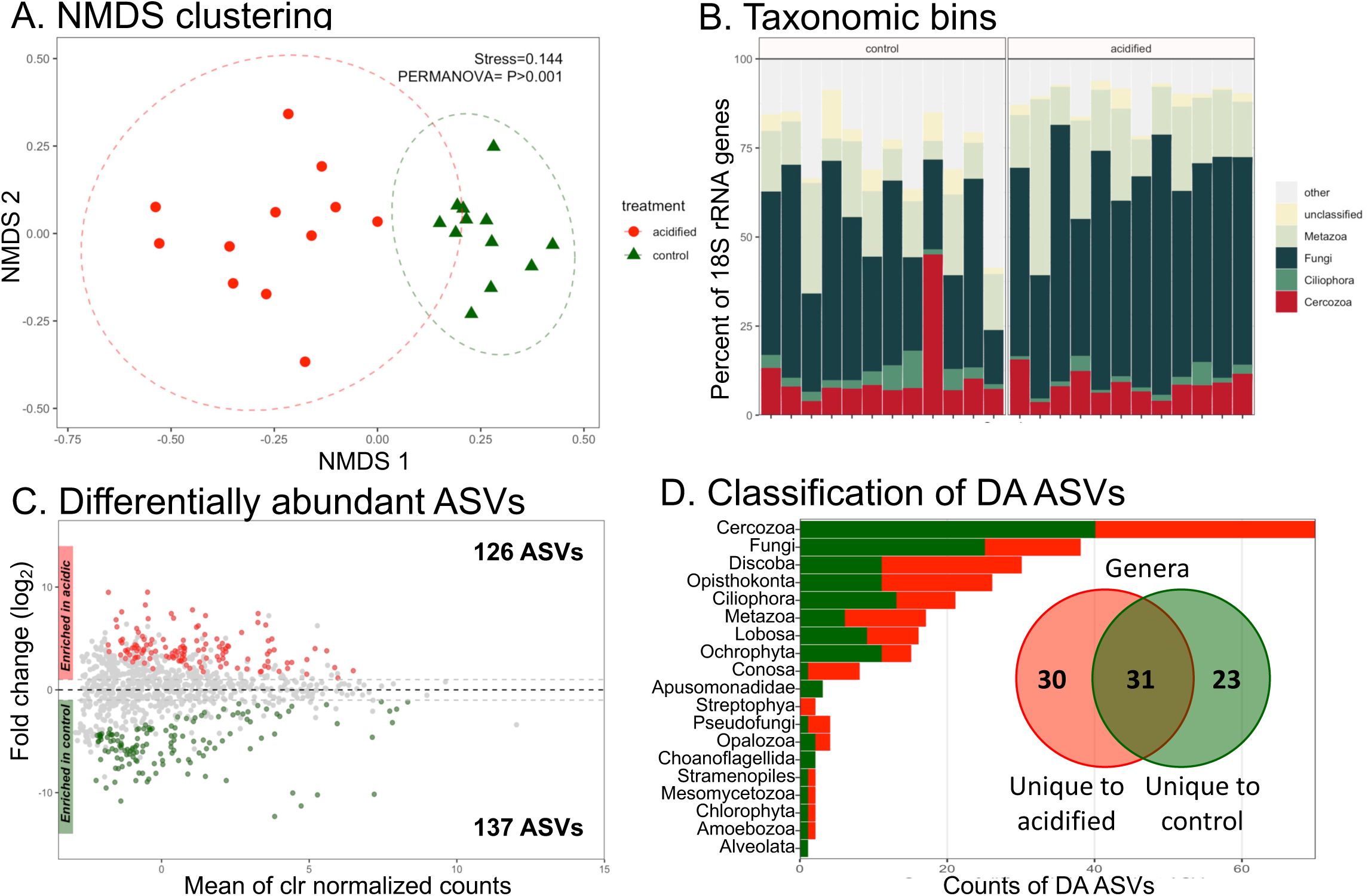
Composition of 18S rRNA gene datasets. **A.** Nonmetric multidimensional scaling ordination on ASV level data. The stress value of the ordination and P-value of a PERMANOVA statistical test are indicated. Ellipses denote 95% confidence intervals fitted onto the ordination. **B. T**axonomic bins in the datasets. Taxa consistently accounting for greater than 1% of sequence reads are displayed with the remainder assigned to the category “other”. A table of taxa showing statistically significant differences in relative abundance are shown in Table S1. **C.** MA plot displaying differentially abundant (DA) ASVs. The number of DA ASVs enriched in each condition are indicated in the inset text. **D.** Taxonomic classification of DA ASVs. Each bar represents the sum of DA ASVs classified to each taxa. The inset Venn diagram shows ASVs classified to the genus level. The diagram shows the sum of genus level bins uniquely enriched in the control or acidified soils, with the overlap indicating genera with members enriched under both conditions.

### Alterations in 18S rRNA libraries due to soil acidification

Assigning 18S rRNA genes to ASVs and visualizing sample relatedness by NMDS demonstrated a significant independent clustering of the control and acidified datasets (P<0.001; Figure 4A). However, contrary to the 16S rRNA datasets the average pairwise distance between control and acidified samples was not significantly different (Figure S1). This suggests that heterogeneity between populations was similar between the control and acidified rhizospheres for the Eukaryotes.

The 18S rRNA genes were classified to explore the relative abundance of taxa in the datasets (Figure 4B). There were wide variations in the average relative abundance of 18S rRNA taxonomic bins between the control and acidified soil datasets. For example, the percent of sequences related to the Fungi increased from an average of 39% of control samples to 56% in the acidified soils, yet the differences were not significant. In fact, only a single taxonomic bin, the Conosa (a subphylum of the Amoebozoa) was identified as significantly different between the control and acidified soils, being more abundant in the control soils (Table S1).

A multitude of 18S rRNA ASVs were identified as DA due to the acidification treatment (Figure 4C). The DA ASVs belonged to 15 different taxonomic ranks, with 4 unclassified bins (Figure 4D). The DA ASVs could be further classified to 84 genus level bins (Figure 4D, inset Venn diagram). One observation of note was that the largest proportion of DA ASVs belonged to the group Cercozoa (Figure 4D), although they were not particularly abundant across the datasets (Figure 4B). This suggests these organisms may be particularly sensitive to the acidification treatment among the Eukaryotes. Yet, the Cercozoa often act as bacterivorous predators, with different feeding strategies (44). In this regard, it is unclear if the Cercozoa are responding to the soil acidification *per se* or an alteration of their prey bacterial populations with acidification. In any case, the data shown here also supports that a wide array of the Eukaryotic rhizosphere communities were sensitive to changes in soil pH, whether this was a response to environmental conditions or changes in the plant health status or bacterial communities remains to be demonstrated.

### Network analysis

Co-occurrence networks were constructed to characterize bacteria, archaea, and eukaryotic interactions. To focus on the most abundant ASVs, only those ASVs with sequence counts greater or equal to 50 and present in both control and acidified datasets were retained. The resulting dataset consisted of 317 16S rRNA gene ASVs and 79 18S rRNA gene ASVs. The co-occurrence network is diagrammed in Figure 5A. Qualitative differences in the network structure are readily apparent from inspecting the network structure. Quantitatively, the Adjusted Rand index between the two networks was 0.016 (two-tailed t-test P<0.001), suggesting that there were highly significant differences in the network topology between the control and acidified soils. Similarly, measurements of network degree (number of connections), eigenvalue centrality (connectedness of nodes) and the hub taxa (taxa that are most connected to other taxa, similar to “keystone” species) all significantly differed between the two networks (Figure 5). Finally, the most central hub taxa were identified for each network (Figure 5B). All of the 5 most central taxa in the control network were bacteria, whereas two Eukaryotes are present in the central taxa of the acidified soil network. In this regard, these data show that the soil acidification treatment did not only alter the structure of the soil community, it also altered how taxa interact and the keystone species that support the community.

**Figure 5.**
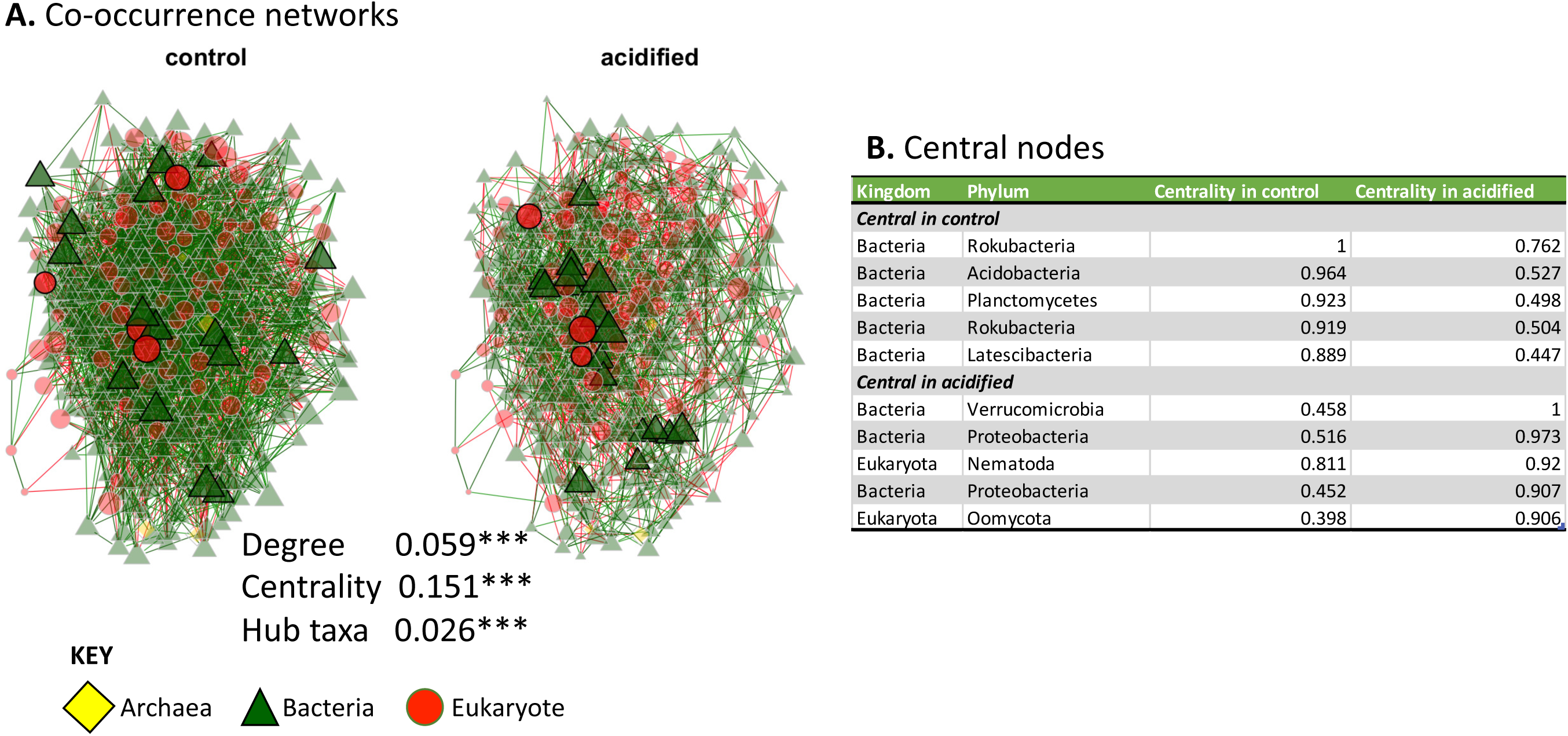
Network analysis. **A.** Each node represents an ASV colored and shaped by the Kingdom to which it belongs. Network connections are colored by their association direction. Positive associations (green) and negative associations (red). The size of the nodes denotes the centrality of the taxon, with larger nodes having the most connections. The absolute difference in degree, eigenvector centrality, and hub taxa measures between the two networks are indicated. Each value was statistically significantly different with a P-value ≤ 0.001 (indicated by the asterisks). **B.** List of the top five most central nodes in the control and acidified soils. Each line represents an individual ASV. The Kingdom and Phylum level classification of each ASV is indicated. The eigenvector centrality which was used to rank the centrality of the ASVs is also displayed.

### Metagenomic sequencing and functional potential of the rhizosphere communities

Bulk sequencing of metagenomic DNA was undertaken to describe the functional gene repertoire of the fir tree rhizosphere communities. Assembly and binning of the reads produced 17 bins from the control metagenomes and 25 bins from the acidified soils (Table S2). The average length of the metagenomic bins was 5.4 Mbp with an average completeness and contamination of 27% and 7%, respectively. None of the assemblies meet the suggested qualifications of even a medium-quality metagenome assembled genome (>90% completion and contamination <5% (45)). Furthermore, a majority of the bins could only be classified to broad taxonomic ranks, with 6 bins being classified to “root” and a further 26 classified to the level of Bacteria (Table S2). Thus, for the remainder of the metagenome analyses the focus is on the genes encoded across the metagenomes rather than focusing on particular Bins.

A total of 127,547 and 157,542 genes were identified in the control and acidified metagenomes, respectively. Of these genes 25.8% and 24.4% were functionally annotated. The KEGG annotations were assigned to pathway modules and a total of 109 and 118 complete modules were identified in the control and acidified metagenomes. Investigating modules that were complete in one treatment but incomplete or absent in the other identified several biochemical pathways that differentiated the control and acidified soils (Table 2). For example, the control metagenomes encoded biosynthesis pathways for isoleucine and thiamine whereas the acidified metagenomes encoded salvage pathways (methionine and thiamine, Table 2). This suggests that the organisms in the control soils may be largely capable of *de-novo* synthesis of amino acids and nucleotides. In contrast, the acidified metagenomes point to auxotrophic pathways requiring amino acid and nucleotide recycling. Another pattern that differentiated the metagenomes was the terminal oxidases identified. Cytochromes of the Cytochrome *c* oxidase, cbb3-type and Cytochrome *o* ubiquinol oxidase in the control soils and Cytochrome *bd* ubiquinol oxidase in the acidified soils. The terminal oxidases in bacteria are regulated based on environmental conditions and can be differentially expressed due to oxygen status, pH, and available electron acceptors (46). Thus, the presence of different cytochromes in the metagenomes between treatments points to adaptations of the microbial communities to inhabiting the control and acidified soils. Finally, the metagenomes from the acidified soils encoded pathways for assimilatory nitrate reduction, indicating a conversion of nitrates to ammonia (47), indicating pathways for nitrogen cycling may have also differed between control and treatment soils. It should be noted that these metagenomes were not exhaustively sampled, so the absence of these pathways in one treatment or the other cannot be taken for an absolute lack of that pathway. Instead, these data suggest that these pathways are not present amongst the most abundant organisms that inhabit the soils from the two treatments.

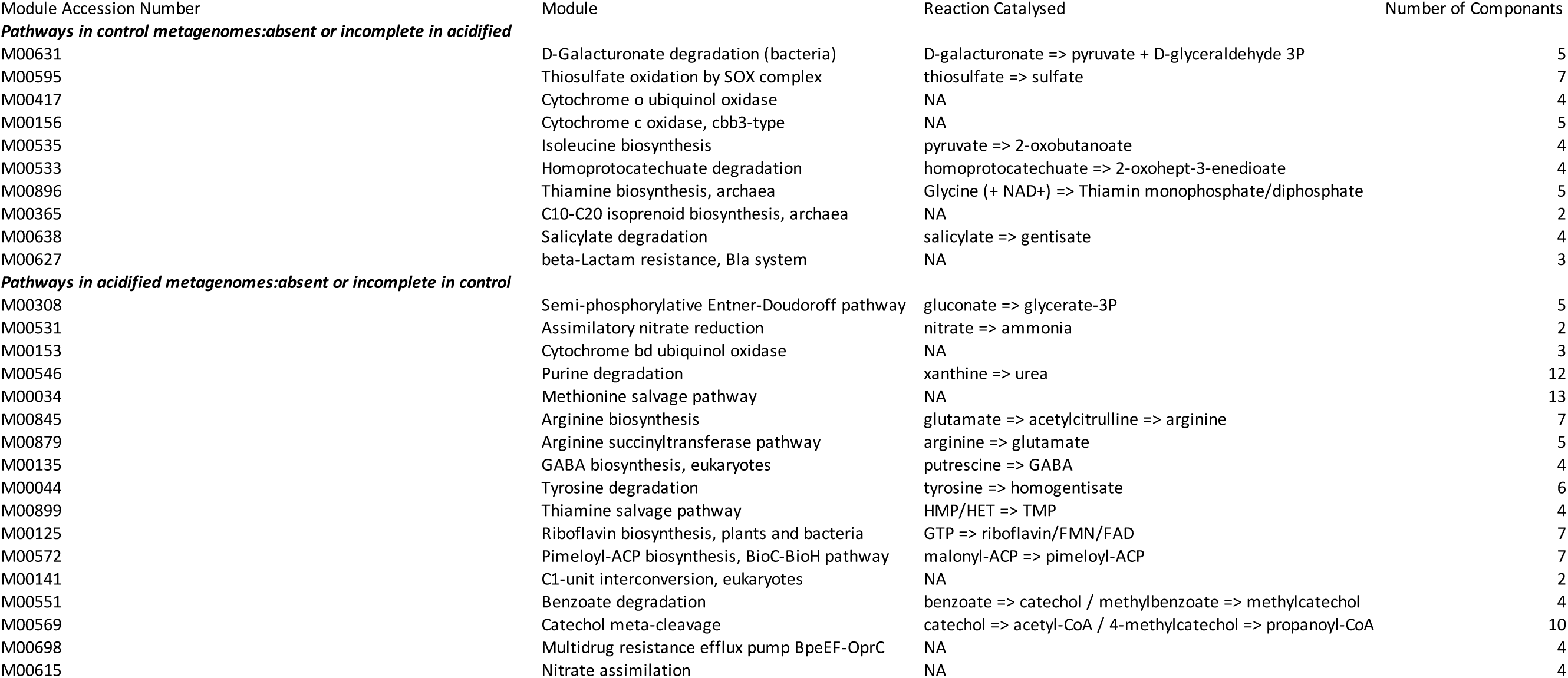

| Module Accession Number | Module | Reaction Catalysed | Number of Components |
| --- | --- | --- | --- |
| ***Pathways in control metagenomes:absent or incomplete in acidified*** |  |  |  |
| M00631 | D-Galacturonate degradation (bacteria) | D-galacturonate => pyruvate + D-glyceraldehyde 3P | 5 |
| M00595 | Thiosulfate oxidation by SOX complex | thiosulfate => sulfate | 7 |
| M00417 | Cytochrome o ubiquinol oxidase | NA | 4 |
| M00156 | Cytochrome c oxidase, cbb3-type | NA | 5 |
| M00535 | Isoleucine biosynthesis | pyruvate => 2-oxobutanoate | 4 |
| M00533 | Homoprotocatechuate degradation | homoprotocatechuate => 2-oxohept-3-enedioate | 4 |
| M00896 | Thiamine biosynthesis, archaea | Glycine (+ NAD+) => Thiamin monophosphate/diphosphate | 5 |
| M00365 | C10-C20 isoprenoid biosynthesis, archaea | NA | 2 |
| M00638 | Salicylate degradation | salicylate => gentisate | 4 |
| M00627 | beta-Lactam resistance, Bla system | NA | 3 |
| ***Pathways in acidified metagenomes:absent or incomplete in control*** |  |  |  |
| M00308 | Semi-phosphorylative Entner-Doudoroff pathway | gluconate => glycerate-3P | 5 |
| M00531 | Assimilatory nitrate reduction | nitrate => ammonia | 2 |
| M00153 | Cytochrome bd ubiquinol oxidase | NA | 3 |
| M00546 | Purine degradation | xanthine => urea | 12 |
| M00034 | Methionine salvage pathway | NA | 13 |
| M00845 | Arginine biosynthesis | glutamate => acetylcitrulline => arginine | 7 |
| M00879 | Arginine succinyltransferase pathway | arginine => glutamate | 5 |
| M00135 | GABA biosynthesis, eukaryotes | putrescine => GABA | 4 |
| M00044 | Tyrosine degradation | tyrosine => homogentisate | 6 |
| M00899 | Thiamine salvage pathway | HMP/HET => TMP | 4 |
| M00125 | Riboflavin biosynthesis, plants and bacteria | GTP => riboflavin/FMN/FAD | 7 |
| M00572 | Pimeloyl-ACP biosynthesis, BioC-BioH pathway | malonyl-ACP => pimeloyl-ACP | 7 |
| M00141 | C1-unit interconversion, eukaryotes | NA | 2 |
| M00551 | Benzoate degradation | benzoate => catechol / methylbenzoate => methylcatechol | 4 |
| M00569 | Catechol meta-cleavage | catechol => acetyl-CoA / 4-methylcatechol => propanoyl-CoA | 10 |
| M00698 | Multidrug resistance efflux pump BpeEF-OprC | NA | 4 |
| M00615 | Nitrate assimilation | NA | 4 |

## DISCUSSION

Soil pH has been identified as a master variable controlling nutrient availability and plant productivity in agricultural soils (48). Here we observe that the soil acidification treatment led to altered mineral nutrient status for the fir tree roots. Specifically, the concentration of Al, Mn, and Zn in the root tissue were significantly higher in root tissues from the acidified soils in comparison to the controls (Figure1). Al is generally abundant in soil but is not considered to be an essential nutrient for plants, and even micromolar amounts of Al in root tissue can be toxic (49). Al toxicity can alter root morphology, lead to deficiencies in other nutrients such as Ca, and cause genetic damage through interactions with plant cellular DNA (reviewed in 38). In comparison, Mn and Zn are required components of the photosynthesis proteins in plants and are thus considered essential nutrients (51). Yet, both nutrients also show toxic effects when provided in excess (52). Thus, these data point to potential stressors in the root tissue related to metal toxicity. In contrast, the levels of B and Ca were reduced in root tissues (Figure 1). B and Ca both play roles in plant cell wall synthesis, and deficiencies are associated with poor plant health (53–55). It is important to note that the trees in the field, including those in the acidified soils, did not show any observable symptoms of phytotoxicity or nutrient deficiency. Indeed, the Canaan fir in this trial is a species adapted to growing in very acid soil; the trees in the acidified plots have shown better growth and color than those growing at a higher soil pH (19). Yet, these observed differences in root mineral nutrient status in association with soil acidification are likely to alter root physiology which presumably could translate into the alterations of the root-associated microbial community in the rhizosphere.

Multiple lines of evidence support that the rhizosphere communities were significantly and dramatically altered in association with soil acidification. Lower soil pH was associated with reductions in alpha-diversity of both the 16S rRNA and 18S rRNA gene datasets, indicating both the bacterial/archaeal and eukaryotic populations were less diverse under soil acidification. Although, the effect and significance were greater for the bacterial/archaeal 16S rRNA gene datasets than for the 18S rRNA genes (Figure 2). Multiple studies have reported a reduction of alpha diversity for microbial populations under soil acidification (56). Thus, a reduction in diversity under soil acidification appears to be a common phenomenon across multiple soil environments.

NMDS clustering, differential abundance of taxonomic bins and ASVs (Figures 3&4), and network analysis (Figure 5) all pointed to an altered microbial community structure in the rhizospheres from control soils in comparison to their acidified counterparts. While other studies have identified particular taxa, such as the Acidobacteria and Actinobacteria as being particularly responsive to soil acidification (18, 57), the data presented here suggests a more generalized response across the community. There was not a strong taxonomic signal in the differentially abundant organisms identified. In the 16S rRNA datasets, multiple phylum-level taxonomic bins and thousands of ASVs belonging to hundreds of individual genera shifted in abundance due to decreased pH (Figure 3). Similar patterns were observed in the 18S rRNA datasets (Figure 4). This broad taxonomic response is likely related to the observation of increased heterogeneity in the rhizosphere communities in the acidified soils, particularly for the 16S rRNA gene datasets (Figure 3A & Figure S1). It has long been recognized that the “coefficient of variation” is a useful tool to assess the stability of ecological communities, where increased variability is a signal of reduced ecological stability (58). In this respect, we propose that the acidified soils may not yet have converged on a stable state, even 6 years after disturbance. Alternatively, the addition of pelletized sulfur to the soil may have simply increased the environmental heterogeneity of the soil, resulting in pH hot spots and cold spots, translating into an elevated inter-sample divergence. Of course, these hypotheses are not mutually exclusive. Yet, taken together these data suggest that a large effect of the soil acidification was a generalized increase in the heterogeneity among the rhizosphere communities, including archaea, bacteria, and eukaryotes, rather than a targeted enrichment or depletion of specific populations.

The large shifts in the taxonomic composition of the soil communities were accompanied by several biochemical pathways that differentiated the functional potential of the sequences recovered from metagenome sequencing of the control and acidified soils (Table 2). For instance, soil acidification was associated with an alteration in terminal oxidases of the respiratory chain encoded in the metagenomes. Genomes from the control soils encoded cytochromes from the cytochrome *c* and cytochrome *o* families, whereas the acidified genomes possessed the genes for cytochrome *bd* (Table 2). Cytochromes of the *bd* family are induced under conditions that are often stressful, such as low O_2_ concentrations (59). In *Escherichia coli* cytochrome *bd* is involved in preventing respiratory inhibition by hydrogen sulfide (60). Given that the method of soil acidification was through addition of sulfur and its interconversion to hydrogen sulfide and sulfuric acid, the presence of cytochrome *bd* may have been protective for the microbes in the acidified soils. This is not the only link to sulfur cycling observed in the metagenomes. Thiosulfate oxidation via the SOX complex was identified as a functional pathway in the control soils but was missing in the acidified soils. Previous studies investigating elemental sulfur cycling in soils identified that *soxB* genes are reduced in abundance and diversity with acidification (61). Thus, these metagenomic data support that the acidification of the soil via elemental sulfur did appear to affect sulfur cycling in the fir tree rhizosphere communities. Additionally, enrichment of cytochrome *bd* is also associated with resistance to nitrosative stress, i.e. the stress induced by excess NO (62). NO is a potent inhibitor of terminal oxidases of the *c* and *o* families as well as inducing oxidative stress (63). Consequently high concentrations of NO in the environment induce a suite of physiological responses. Nitric oxide (NO) is a product of the reduction of nitrate to ammonia (64), a pathway specifically encoded in the metagenomes in the acidified soils (Table 2) and thus, indicating a potential for nitrosative stress in the rhizosphere soils. Taken together, these metagenomic data point to an alteration in the environmental conditions and nutrient cycling in the rhizosphere community.

## Conclusion

The data presented here demonstrates that the effects of soil acidification are manifest across the range of organisms that inhabit the soil. This includes changes in the mineral nutrient status of the host plant as well as compositional alterations in the associated archaeal, bacterial, and eukaryotic communities that populate the rhizosphere soils. The microbial communities were less diverse in the acidified soils and alterations in the taxonomic composition of the communities were evident at multiple taxonomic ranks, suggesting that a wide variety of the soil microbial community were affected by the acidification treatment. The results of these taxonomic shifts resulted in an increase in the heterogeneity in community structure amongst the microbial populations in the acidified soils. This could indicate that the acidified soils are in a transitional state or inhabit an environment with increased spatial heterogeneity. Finally, metagenome sequencing demonstrated that the taxonomic reshaping of the community translated into alterations in the functional potential of the indigenous rhizosphere populations. Pathways involved in carbon, sulfur, and nitrogen cycling were differentially present between the metagenomes from control and acidified soils, linking the changes in the taxonomic composition of the communities to their functional potential and nutrient cycling. These data underscore the importance of soil pH as a driving force in determining the structure and function of soil communities and highlights the critical research need to integrate plant and microbial responses in the rhizosphere and their responses to soil acidification.

## Data accessibility

All 16S rRNA, 18S rRNA amplicon gene libraries, and the shotgun metagenome sequences are available in the NCBI SRA under the BioProject accession number PRJNA708254. The *Abies balsamea* var. phanerolepis 18S rRNA gene sequence is available in Genbank under the accession number MW699166.1.

## Acknowledgments

We thank the Christmas Tree Promotion Board for funding “Investigating soil acidification mechanisms for inhibiting *Phytophthora”.* ST was supported by an AFRI Foundational Program grant from the United States Department of Agriculture–National Institute of Food and Agriculture (USDA-NIFA grant number 2019-67019-29315). This work was additionally supported by the USDA National Institute of Food and Agriculture, Hatch project 1022006 awarded to BS, and Hatch project 1012247 awarded to RSC.

## References

1. Soil Survey Staff. 2014. Soil Survey Field and Laboratory Methods Manual. Soil Survey Investigations Report No. 51, Version 2.0. U.S. Department of Agriculture, Natural Resources Conservation Service.

2. Kunhikrishnan A, Thangarajan R, Bolan NS, Xu Y, Mandal S, Gleeson DB, Seshadri B, Zaman M, Barton L, Tang C, Luo J, Dalal R, Ding W, Kirkham MB, Naidu R. 2016. Functional Relationships of Soil Acidification, Liming, and Greenhouse Gas Flux, p. 1–71. In Advances in Agronomy. Elsevier.

3. Azevedo LB, van Zelm R, Hendriks AJ, Bobbink R, Huijbregts MAJ. 2013. Global assessment of the effects of terrestrial acidification on plant species richness. Environmental Pollution 174:10–15.

4. Raza S, Miao N, Wang P, Ju X, Chen Z, Zhou J, Kuzyakov Y. 2020. Dramatic loss of inorganic carbon by nitrogen-induced soil acidification in Chinese croplands. Global Change Biology 26:3738–3751.

5. Meng C, Tian D, Zeng H, Li Z, Yi C, Niu S. 2019. Global soil acidification impacts on belowground processes. Environ Res Lett 14:074003.

6. Haynes RJ. 1983. Soil acidification induced by leguminous crops. Grass and Forage Science 38:1–11.

7. Barak P, Jobe BO, Krueger AR, Peterson LA, Laird DA. 1997. Effects of long-term soil acidification due to nitrogen fertilizer inputs in Wisconsin. Plant and Soil 197:61–69.

8. Bolan NS, Hedley MJ. 2003. Role of carbon, nitrogen, and sulfur cycles in soil acidification. Handbook of soil acidity 29–56.

9. Evans CD, Jones TG, Burden A, Ostle N, Zieliński P, Cooper MDA, Peacock M, Clark JM, Oulehle F, Cooper D, Freeman C. 2012. Acidity controls on dissolved organic carbon mobility in organic soils. Global Change Biology 18:3317–3331.

10. Choma M, Tahovská K, Kaštovská E, Bárta J, Růžek M, Oulehle F. 2020. Bacteria but not fungi respond to soil acidification rapidly and consistently in both a spruce and beech forest. FEMS Microbiology Ecology 96.

11. Ekström SM, Kritzberg ES, Kleja DB, Larsson N, Nilsson PA, Graneli W, Bergkvist B. 2011. Effect of Acid Deposition on Quantity and Quality of Dissolved Organic Matter in Soil–Water. Environ Sci Technol 45:4733–4739.

12. Fierer N, Jackson RB. 2006. The diversity and biogeography of soil bacterial communities. PNAS 103:626–631.

13. Fierer N. 2017. Embracing the unknown: disentangling the complexities of the soil microbiome. Nature Reviews Microbiology 15:579–590.

14. Hartman WH, Richardson CJ, Vilgalys R, Bruland GL. 2008. Environmental and anthropogenic controls over bacterial communities in wetland soils. PNAS 105:17842– 17847.

15. Lauber CL, Hamady M, Knight R, Fierer N. 2009. Pyrosequencing-Based Assessment of Soil pH as a Predictor of Soil Bacterial Community Structure at the Continental Scale. Appl Environ Microbiol 75:5111–5120.

16. Nye PH. 1981. Changes of pH across the rhizosphere induced by roots. Plant Soil 61:7–26.

17. Wang X, Tang C, Severi J, Butterly CR, Baldock JA. 2016. Rhizosphere priming effect on soil organic carbon decomposition under plant species differing in soil acidification and root exudation. New Phytologist 211:864–873.

18. Shen G, Zhang S, Liu X, Jiang Q, Ding W. 2018. Soil acidification amendments change the rhizosphere bacterial community of tobacco in a bacterial wilt affected field. Appl Microbiol Biotechnol 102:9781–9791.

19. Cowles RS. 2020. Sulfur Amendment of Soil Improves Establishment and Growth of Firs in a Field Naturally Infested with Phytophthora1. Journal of Environmental Horticulture 38:15– 21.

20. McPherson MR, Wang P, Marsh EL, Mitchell RB, Schachtman DP. 2018. Isolation and Analysis of Microbial Communities in Soil, Rhizosphere, and Roots in Perennial Grass Experiments. JoVE (Journal of Visualized Experiments) e57932.

21. Alivisatos AP, Blaser MJ, Brodie EL, Chun M, Dangl JL, Donohue TJ, Dorrestein PC, Gilbert JA, Green JL, Jansson JK, Knight R, Maxon ME, McFall-Ngai MJ, Miller JF, Pollard KS, Ruby EG, Taha SA, Unified Microbiome Initiative Consortium. 2015. A unified initiative to harness Earth’s microbiomes. Science 350:507–508.

22. Lundberg DS, Yourstone S, Mieczkowski P, Jones CD, Dangl JL. 2013. Practical innovations for high-throughput amplicon sequencing. Nature Methods 10:999–1002.

23. Taerum SJ, Steven B, Gage DJ, Triplett LR. 2020. Validation of a PNA Clamping Method for Reducing Host DNA Amplification and Increasing Eukaryotic Diversity in Rhizosphere Microbiome Studies. Phytobiomes Journal 4:291–302.

24. Medlin L, Elwood HJ, Stickel S, Sogin ML. 1988. The characterization of enzymatically amplified eukaryotic 16S-like rRNA-coding regions. Gene 71:491–499.

25. Quast C, Pruesse E, Yilmaz P, Gerken J, Schweer T, Yarza P, Peplies J, Glöckner FO. 2012. The SILVA ribosomal RNA gene database project: improved data processing and web-based tools. Nucl Acids Res D590–6.

26. Schloss PD, Westcott SL, Ryabin T, Hall JR, Hartmann M, Hollister EB, Lesniewski RA, Oakley BB, Parks DH, Robinson CJ, Sahl JW, Stres B, Thallinger GG, Horn DJV, Weber CF. 2009. Introducing mothur: Open-source, platform-independent, community-supported software for describing and comparing microbial communities. Appl Environ Microbiol 75:7537–7541.

27. Rognes T, Flouri T, Nichols B, Quince C, Mahé F. 2016. VSEARCH: a versatile open source tool for metagenomics. PeerJ 4:e2584.

28. Guillou L, Bachar D, Audic S, Bass D, Berney C, Bittner L, Boutte C, Burgaud G, de Vargas C, Decelle J, del Campo J, Dolan JR, Dunthorn M, Edvardsen B, Holzmann M, Kooistra WHCF, Lara E, Le Bescot N, Logares R, Mahé F, Massana R, Montresor M, Morard R, Not F, Pawlowski J, Probert I, Sauvadet A-L, Siano R, Stoeck T, Vaulot D, Zimmermann P, Christen R. 2013. The Protist Ribosomal Reference database (PR2): a catalog of unicellular eukaryote Small Sub-Unit rRNA sequences with curated taxonomy. Nucleic Acids Res 41:D597–D604.

29. Cole JR, Chai B, Farris RJ, Wang Q, Kulam SA, McGarrell DM, Garrity GM, Tiedje JM. 2005. The Ribosomal Database Project (RDP-II): sequences and tools for high-throughput rRNA analysis. Nucleic Acids Res 33:D294–D296.

30. McMurdie PJ, Holmes S. 2013. phyloseq: an R package for reproducible interactive analysis and graphics of microbiome census data. PloS one 8:e61217.

31. Fernandes AD, Macklaim JM, Linn TG, Reid G, Gloor GB. 2013. ANOVA-Like Differential Expression (ALDEx) Analysis for Mixed Population RNA-Seq. PLoS ONE 8:e67019.

32. Peschel S, Müller CL, von Mutius E, Boulesteix A-L, Depner M. 2020. NetCoMi: network construction and comparison for microbiome data in R. Briefings in Bioinformatics https://doi.org/10.1093/bib/bbaa290.

33. Kurtz ZD, Müller CL, Miraldi ER, Littman DR, Blaser MJ, Bonneau RA. 2015. Sparse and Compositionally Robust Inference of Microbial Ecological Networks. PLOS Computational Biology 11:e1004226.

34. Yoon G, Gaynanova I, Müller CL. 2019. Microbial Networks in SPRING - Semi-parametric Rank-Based Correlation and Partial Correlation Estimation for Quantitative Microbiome Data. Front Genet 10.

35. Kolmogorov M, Yuan J, Lin Y, Pevzner PA. 2019. Assembly of long, error-prone reads using repeat graphs. 5. Nature Biotechnology 37:540–546.

36. Kolmogorov M, Bickhart DM, Behsaz B, Gurevich A, Rayko M, Shin SB, Kuhn K, Yuan J, Polevikov E, Smith TPL, Pevzner PA. 2020. metaFlye: scalable long-read metagenome assembly using repeat graphs. 11. Nature Methods 17:1103–1110.

37. Vaser R, Sović I, Nagarajan N, Šikić M. 2017. Fast and accurate de novo genome assembly from long uncorrected reads. Genome Res 27:737–746.

38. Kang DD, Li F, Kirton E, Thomas A, Egan R, An H, Wang Z. 2019. MetaBAT 2: an adaptive binning algorithm for robust and efficient genome reconstruction from metagenome assemblies. PeerJ 7.

39. Parks DH, Imelfort M, Skennerton CT, Hugenholtz P, Tyson GW. 2015. CheckM: assessing the quality of microbial genomes recovered from isolates, single cells, and metagenomes. Genome Res 25:1043–1055.

40. Hyatt D, Chen G-L, LoCascio PF, Land ML, Larimer FW, Hauser LJ. 2010. Prodigal: prokaryotic gene recognition and translation initiation site identification. BMC Bioinformatics 11:119.

41. Suzuki S, Ishida T, Ohue M, Kakuta M, Akiyama Y. 2017. GHOSTX: A Fast Sequence Homology Search Tool for Functional Annotation of Metagenomic Data, p. 15–25. *In* Kihara, D (ed.), Protein Function Prediction: Methods and Protocols. Springer, New York, NY.

42. Suzuki S, Kakuta M, Ishida T, Akiyama Y. 2014. GHOSTX: An Improved Sequence Homology Search Algorithm Using a Query Suffix Array and a Database Suffix Array. PLOS ONE 9:e103833.

43. Kanehisa M, Araki M, Goto S, Hattori M, Hirakawa M, Itoh M, Katayama T, Kawashima S, Okuda S, Tokimatsu T, Yamanishi Y. 2008. KEGG for linking genomes to life and the environment. Nucleic Acids Research 36:D480–D484.

44. Glücksman E, Bell T, Griffiths RI, Bass D. 2010. Closely related protist strains have different grazing impacts on natural bacterial communities. Environmental Microbiology 12:3105– 3113.

45. Bowers RM, Kyrpides NC, Stepanauskas R, Harmon-Smith M, Doud D, Reddy TBK, Schulz F, Jarett J, Rivers AR, Eloe-Fadrosh EA, Tringe SG, Ivanova NN, Copeland A, Clum A, Becraft ED, Malmstrom RR, Birren B, Podar M, Bork P, Weinstock GM, Garrity GM, Dodsworth JA, Yooseph S, Sutton G, Glöckner FO, Gilbert JA, Nelson WC, Hallam SJ, Jungbluth SP, Ettema TJG, Tighe S, Konstantinidis KT, Liu W-T, Baker BJ, Rattei T, Eisen JA, Hedlund B, McMahon KD, Fierer N, Knight R, Finn R, Cochrane G, Karsch-Mizrachi I, Tyson GW, Rinke C, Lapidus A, Meyer F, Yilmaz P, Parks DH, Eren AM, Schriml L, Banfield JF, Hugenholtz P, Woyke T. 2017. Minimum information about a single amplified genome (MISAG) and a metagenome-assembled genome (MIMAG) of bacteria and archaea. 8. Nature Biotechnology 35:725–731.

46. Jurtshuk P, Mueller TJ, Acord WC. 1975. Bacterial Terminal Oxidases. CRC Critical Reviews in Microbiology 3:399–468.

47. Lin JT, Stewart V. 1997. Nitrate Assimilation by Bacteria, p. 1–30. In Poole, RK (ed.), Advances in Microbial Physiology. Academic Press.

48. Msimbira LA, Smith DL. 2020. The Roles of Plant Growth Promoting Microbes in Enhancing Plant Tolerance to Acidity and Alkalinity Stresses. Front Sustain Food Syst 4.

49. Delhaize E, Ryan PR. 1995. Aluminum Toxicity and Tolerance in Plants. Plant Physiol 107:315–321.

50. Gupta N, Gaurav SS, Kumar A. 2013. Molecular Basis of Aluminium Toxicity in Plants: A Review. American Journal of Plant Sciences 2013.

51. Hänsch R, Mendel RR. 2009. Physiological functions of mineral micronutrients (Cu, Zn, Mn, Fe, Ni, Mo, B, Cl). Current Opinion in Plant Biology 12:259–266.

52. Millaleo R, Reyes-Diaz M, Ivanov AG, Mora ML, Alberdi M. 2010. MANGANESE AS ESSENTIAL AND TOXIC ELEMENT FOR PLANTS: TRANSPORT, ACCUMULATION AND RESISTANCE MECHANISMS. Journal of soil science and plant nutrition 10:470–481.

53. Liu Y, Riaz M, Yan L, Zeng Y, Cuncang J. 2019. Boron and calcium deficiency disturbing the growth of trifoliate rootstock seedlings (Poncirus trifoliate L.) by changing root architecture and cell wall. Plant Physiology and Biochemistry 144:345–354.

54. Brdar-Jokanović M. 2020. Boron Toxicity and Deficiency in Agricultural Plants. 4. International Journal of Molecular Sciences 21:1424.

55. Parvin K, Nahar K, Hasanuzzaman M, Bhuyan MHMB, Fujita M. 2019. Calcium-Mediated Growth Regulation and Abiotic Stress Tolerance in Plants, p. 291–331. *In* Hasanuzzaman, M, Hakeem, KR, Nahar, K, Alharby, HF (eds.), Plant Abiotic Stress Tolerance: Agronomic, Molecular and Biotechnological Approaches. Springer International Publishing, Cham.

56. Zhang X, Liu W, Zhang G, Jiang L, Han X. 2015. Mechanisms of soil acidification reducing bacterial diversity. Soil Biology and Biochemistry 81:275–281.

57. Wu Y, Zeng J, Zhu Q, Zhang Z, Lin X. 2017. pH is the primary determinant of the bacterial community structure in agricultural soils impacted by polycyclic aromatic hydrocarbon pollution. 1. Scientific Reports 7:40093.

58. Kéfi S, Domínguez-García V, Donohue I, Fontaine C, Thébault E, Dakos V. 2019. Advancing our understanding of ecological stability. Ecology Letters 22:1349–1356.

59. Borisov VB, Gennis RB, Hemp J, Verkhovsky MI. 2011. The cytochrome bd respiratory oxygen reductases. Biochimica et Biophysica Acta (BBA) - Bioenergetics 1807:1398–1413.

60. Korshunov S, Imlay KRC, Imlay JA. 2016. The cytochrome bd oxidase of Escherichia coli prevents respiratory inhibition by endogenous and exogenous hydrogen sulfide. Molecular Microbiology 101:62–77.

61. Zhao C, Gupta VV, S, R, Degryse F, Mclaughlin MJ. 2017. Effects of pH and ionic strength on elemental sulphur oxidation in soil. Biology and Fertility of Soils 53:247–256.

62. Giuffrè A, Borisov VB, Arese M, Sarti P, Forte E. 2014. Cytochrome bd oxidase and bacterial tolerance to oxidative and nitrosative stress. Biochimica et Biophysica Acta (BBA) - Bioenergetics 1837:1178–1187.

63. Cooper CE. 2002. Nitric oxide and cytochrome oxidase: substrate, inhibitor or effector? Trends in Biochemical Sciences 27:33–39.

64. Cole JA. 2018. Anaerobic Bacterial Response to Nitrosative Stress, p. 193–237. In Advances in Microbial Physiology. Elsevier.

